# Characterization of DNA replicators of a 75-kb region of Chromosome 2 of fission yeast, *Schizosacharomyces pombe*

**DOI:** 10.1101/2022.07.01.498468

**Authors:** Mukesh Pratap Yadav, Dharani Dhar Dubey

## Abstract

DNA replication origins of a 75-kb region of chromosome 2 of *Schizosacharomyces pombe*, are well characterized. Here, we have further characterized two origins, ars2-3358 and ars2-3363 on plasmids using an Autonomous Replicating Sequences (ARS) assay. Sequence analysis of the five ARSs located within the 75-kb region revealed several asymmetric AT motifs ≥10 bp in each of these origins. Interestingly, we have also found a 29 bp AT-rich sequence (29W) in ars2-3358, which alone is not sufficient to provide DNA replication origins to plasmid. A genomic search for asymmetric AT motifs ≥10 bp gave a total 2309 near matches to 9 of these motifs in the genome. Majority (>65% to >77%) of six of these AT motifs overlapped with the earlier mapped genome-wide DNA replication origins. Moreover, functional comparison of these AT motifs to the deletion and mutational studies showed strong correlation with DNA replication origins efficiency. Therefore, we hypothesize that these sequences are important redundant core elements of *S. pombe* origins.

## Introduction

DNA replication maintains genomic integrity and duplicates precisely the genome of an organism during the S phase of the cell division cycle. It is a highly coordinated mechanism of *cis* and *trans* elements. Any disparity in the coordination of these elements may lead to the genetic instability and ultimately disease. In prokaryotes, single DNA replication origin are defined to replicate whole genome [1]. However, in eukaryotes hundreds to thousands of DNA replication origins, required to replicate the larger eukaryotic genomes, are spread along the chromosomes [2].

In the yeasts *Saccharomyces cerevisiae* (*S. cerevisiae*) and *Schizosaccharomyces pombe* (*S. pombe*) origins are well characterized and are referred to as <u>A</u>utonomous <u>R</u>eplicating <u>S</u>equences (ARS). ARSs are small genomic DNA sequences (100-150 bp in *S. cerevisiae* and 500-1500 bp in *S. pombe*), which confer high efficiency autonomous replication on plasmids, and allow for stable plasmid maintenance [3–5]. DNA Sequence analysis of *S. cerevisiae* ARS’s revealed three modular elements: A, B and C [6]. Furthermore, mutational and genomic analysis showed that ARSs contain a 17-bp conserved sequence in a domain A, called as <u>A</u>RS <u>C</u>onsensus <u>S</u>equence (ACS), flanked on one side by nucleosome-depleted stretch of DNA [7,8].

In *S. pombe*, ARSs are less defined than in *S. cerevisiae* [9–15]. They are larger in size (500-2000 bp), highly AT-rich, contain multiple redundant functional domains and do not possess any consensus sequence [10,13,15–17]. Moreover, in *S. pombe* there are multiple redundant sequences, which can be either substituted by other AT-rich sequences or deleted without significantly affecting the ARS activity [10,13,15–17].

Okuno et al. (1999) identified five ARS elements in a 100-kb region of chromosome 2 of *S. pombe*. Three of these ARSs were found to be active as chromosomal origins and their transformation frequencies were roughly proportional to their origin efficiency. Cotobal et al. (2010) reported AT-rich fragments shorter than 100 bp with no homology in the genome of *S. pombe* are capable of initiating DNA replication in the genome. Moreover, functional dissection of endogenous origins revealed several 25-30 bp contiguous A-T motifs in the origins capable of initiating DNA replication.

Our lab mapped five new ARS elements (*ars2006-ars2011*, ars2-3335 through ars2-3398 in the systematic nomenclature (ars2 stand for Autonomous Replicating Sequences in chromosome 2, last four numerical digit represent the known origin) corresponding to five new replication origins [19,20], in another 75-kb stretch of chromosome 2. The correspondence between the chromosomal origins and their plasmid ARS activity were not precise because of the two ARSs were not precisely localized. However, most of the ARSs and origins of 75-kb stretches overlap with genome-wide studies [21–26]. Hence, it was important to precisely localize these ARSs and compare their transformation frequency with chromosomal origin efficiency.

Here, we have precisely localized the ARSs, compared ARS and genomic origin efficiency, and also analyzed DNA sequences of ARSs, nucleosome occupancy and AT-rich motifs. Interestingly, one of the origin, ars2-3358 contains a 29 bp AT-rich sequence (29W) sequence, though it is not sufficient for ARS activity. In addition, several AT-contiguous ≥10 bp motifs were found in each of the five ARSs. We refer to these AT-contiguous motifs as polyAT. Approximately 2300 (Supplementary Table 1) similar sequences were found across the genome in the BLAST. They overlap with ≥65% of the recently mapped genome-wide origins [21].

## Materials and Method

### Strains and Media

For cloning experiments, *Escherichia coli* DH5α cells were grown in Luria-Bertani medium supplemented with 100 μg/ml ampicillin at 37°C; solid media contained 2% agar. The *S. pombe* strain D18 (*ura4-*D18 *leu1-32 h*) was used for transformation assays. For genomic DNA preparation, D18 cells were grown at 30°C in YES supplemented with 150 mg/litre each adenine, leucine and uracil. After transformation, D18 cells were plated onto EMM medium supplemented with leucine and incubated at 30°C.

### Cloning

For cloning, the parental plasmids e.g., ars2-3358 (*ars2009*-4618 bp) and ars2-3363 (*ars2010*-2348 bp) were used. The cloning of three fragments of ars2-3358 were done by restriction digestion of 4618 bp plasmid with following restriction enzymes *Eco*RI-*Bgl*II (2662 bp), *Cla*I-*Pvu*II (2452 bp) and *Bgl*II-*Eco*RI (1956 bp) and 2662 bp and 1956 bp fragment were ligated into the pRS306 digested with *Eco*RI-*Bam*HI and 2452 bp into *Cla*I-*Sma*I. However, promoter deletion of ars2-3363 (*ars2010*) was done by PCR based (primer list, Table 1) cloning strategy. PCR was done using the primers mentioned in Table 1 which contain *Hind*III site. The PCR fragment was purified and digested by *H°ind*III and ligated into *Hind*III digested pRS306. The plasmid pRS306, pBluescript SK+ derivative containing *ura3* gene of *S. cerevisiae*, which complements to *ura4* gene of *S. pombe* was used as cloning vector. All sequences were downloaded from pomBase (https://www.pombase.org). The regions of chromosome 2 from the position 3325424-3400425 were used for analysis of ARSs and origins (Figure 1A).

**Table 1.**
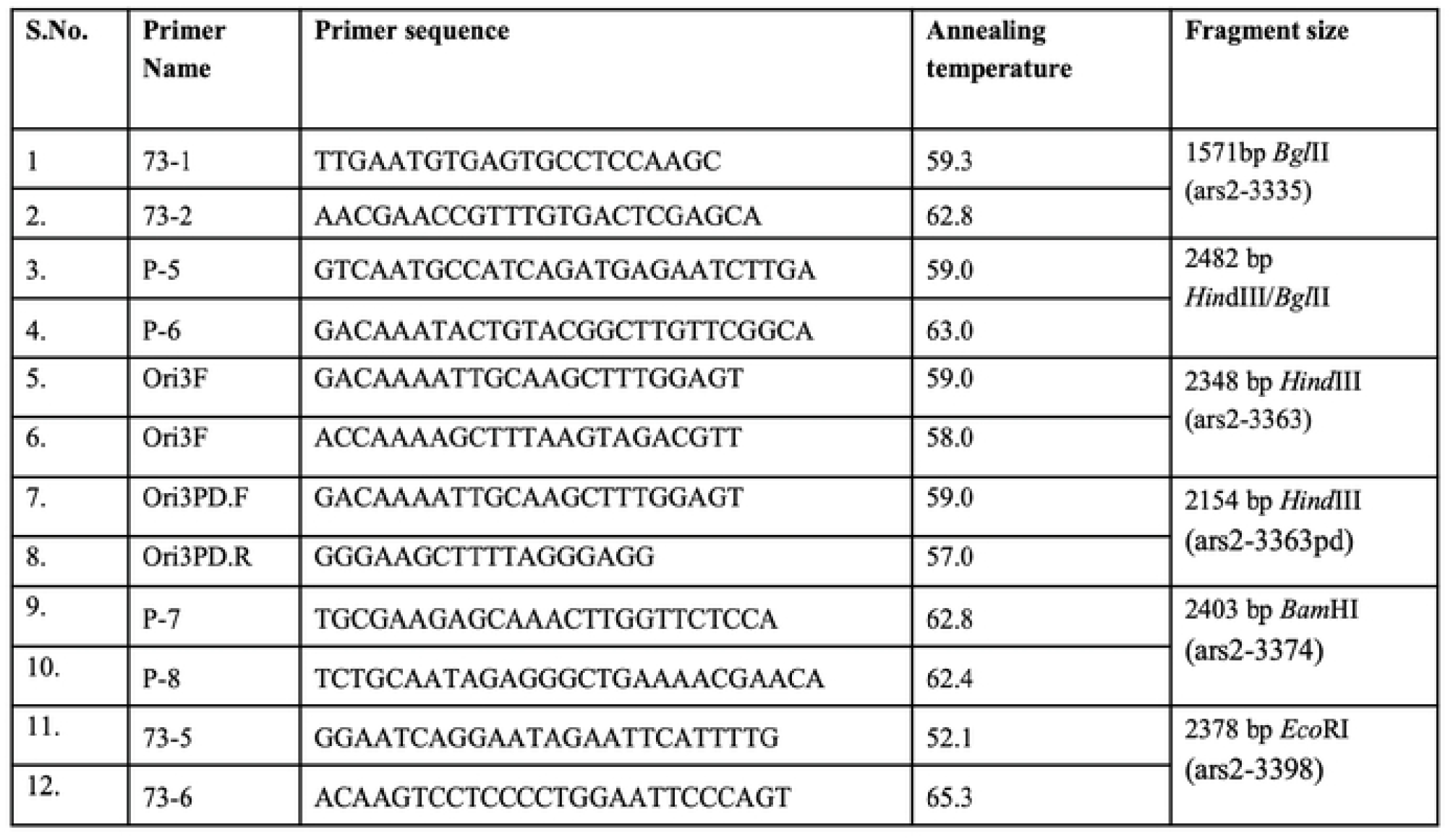
List of primers, primer sequences, annealing temperature and their fragment size along with restriction enzymes.

**Figure 1:**
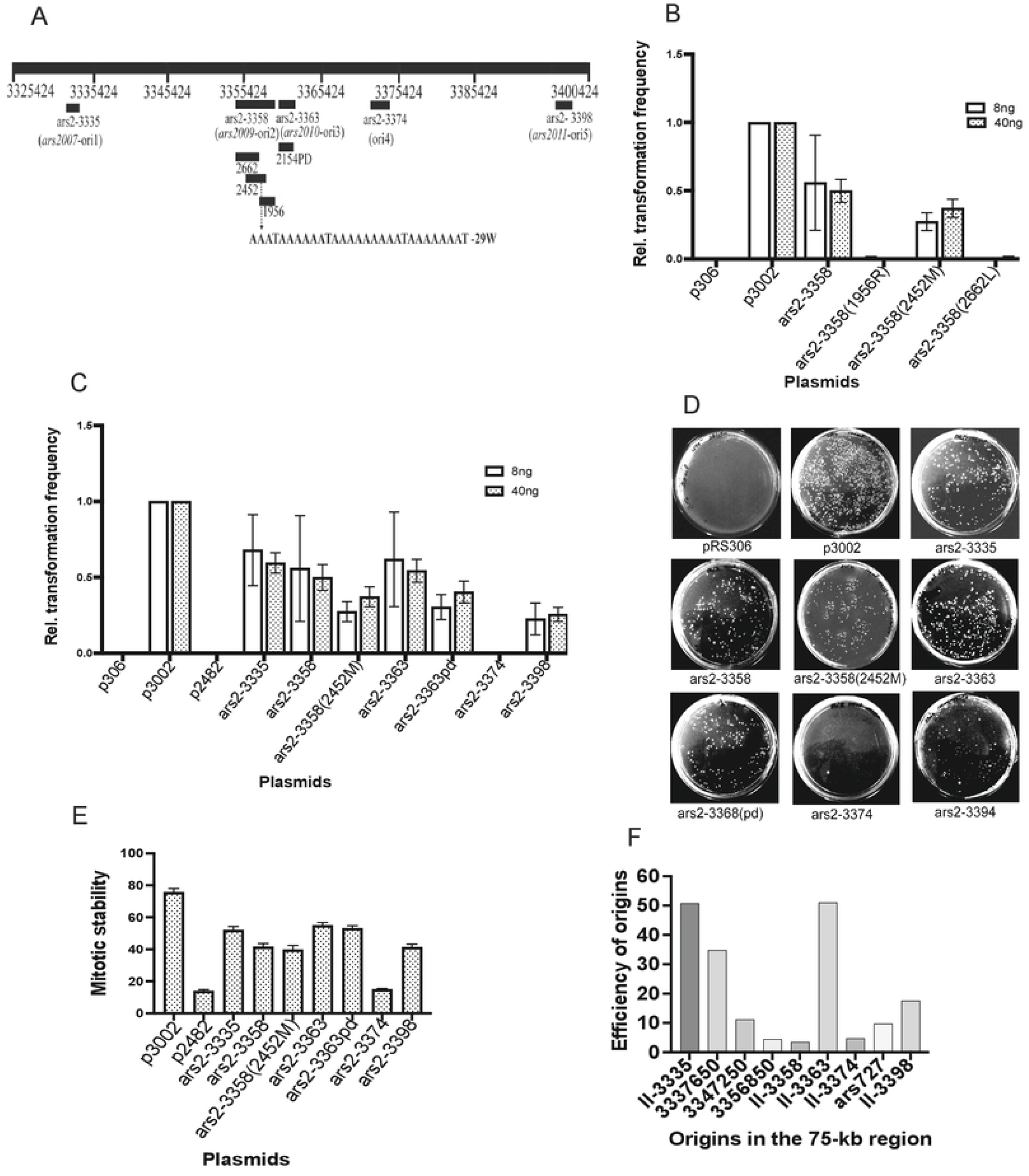
75-kb region of the chromosome 2 showing origins/ARSs mapped by Dubey et al., 2010. Precise location, transformation frequencies and mitotic stability analysed in the study. (A)- Line diagram marked with chromosome positions, origins and ARSs located in the 75-kb region. (B)- Relative transformation frequencies of ars2-3358 and subframent cloned. (C)- Combined relative transformation frequency of precisely localized ARSs of 75-kb region. (D)-Plate photograph of D18 transformed cells plated on EMM uracil negative agar plate transformed with precisely cloned ARSs. (E)- Mitotic stability of the plasmids. (F)- comparison of75-kb region origins efficiency between two studies Daigaku et al. 2015 and Dubey et al. 2010.

### ARS assay

To test ARS activity, a lithium acetate protocol [27] was used to transform D18 *S. pombe* cells. D18 cells were transformed with 8 ng (20μL D18 transformed cells) or 40 ng (100μL D18 transformed cells) of DNA and plated on EMM plates supplemented with leucine for selection. Photographs of the plates with the transformants were taken and the colonies were counted after 5-7 days of incubation at 30°C. p3002 and pRS306 were used as positive and negative controls respectively.

### Mitotic stability and status of plasmid maintenance

To test the mitotic stability and maintenance status of plasmid, experiments were performed as described [28].

### DNA Preparation

The plasmid DNA was isolated using the standard alkaline lysis protocol [29]. For PCR, genomic DNA from 972h^-^ *S. pombe* cells was prepared using the smash and grab method of Hoffman & Winston, (1987).

### Helical stability and Nucleosome occupancy

Stress Induced DNA duplex Destabilization (SIDD) of origin fragments was analyzed with WebSIDD https://benham.genomecenter.ucdavis.edu/sibz [31,32]. Nucleosome occupancy was analyzed using Segal’s lab online tool (http://genie.weizmann.ac.il/software/nucleo_prediction.html) [33]

### Sequence analysis

All the sequences were analyzed using FastPCR, pDraw32, MEME and MS-Word (figures were made using adobe photoshop3, adobe illustrator 2022 and Biorander (https://app.biorender.com/illustrations). Also, the AT-contiguous stretches found were used for blast using pomBase <https://www.pombase.org>, keeping following parameters -

a. - Searching sensitivity= near match
b. - Maximum no. of alignment displayed = 05
c. - Minimum no. of score displayed = 05
d. - E-value threshold = 1.0
e. - Filter low complexity regions = no.

## Results and Discussion

### Precise location and strength testing of ARSs

We have mapped five chromosomal replication origins in a 75-kb stretch of chromosome 2 (Figure 1A) and estimated their efficiency using two dimensional agarose gel electrophoresis [20]. Three of these origins, II-3335, II-3363 and II-3398 are moderately active and two, II-3358 and II-3374, are weak. The DNA fragments encompassing each of these origins were cloned in pRS306 and tested for their origin activity in isolation (free from any influence of the chromosomal context including nearby origins) by ARS and mitotic stability-assay. A combination of these two assays provides a robust test for origin activity as the plasmids with efficient origin sequences transform yeast cells at a high frequency, show high mitotic stability and are maintained mostly as monomers. The plasmids containing II-3335 (ars2-3335), II-3358 (ars2-3358), II-3363 (ars2-3358) and II-3398 (ars2-3398) were ARS positive while the one with II-3374 (ars2-3374) was ARS negative. Although II-3358 was a weak chromosomal origin, the ARS activity of ars2-3358 was equal to the plasmids derived from the other moderately active origins (II-3363,[20]).

In this study, we modified two of these origin sequences as described below and used the resulting plasmids along with the other origin-containing plasmids for ARS and mitotic stability assays. The 4618 bp, II-3358 fragment was sub cloned as three overlapping fragments of 2662 bp (ars2-3358(2662)), 2452 bp (ars2-3358(2452M)) and 1956 bp (ars2-3358(1956)) to precisely localize the sequences important for ARS activity. The results of transformation (Figure 1B) and mitotic stability experiments (Figure 1E) show the presence of ARS activity in ars2-3358(2452M) with slightly reduced efficiency than the parental plasmid ars2-3358. The other two plasmids are ARS negative. Noteworthy, the 29W sequence reported to be important for origin activity [18] is present in both ars2-3358(2452M) (close to its right end) and ars2-3358(1956) (near its left end). The lack of ARS activity in ars2-3358(1956) suggests that the 29W sequence alone is not sufficient for the ARS activity. Additional sequences present along with the 29W in ars2-3358(2452M) are required for ARS activity, an observation similar to that of Cotobal et al. (2010). The reduction in ARS efficiency of ars2-3358(2452M) could be due to the size reduction of the cloned fragment and might represent the presence of auxiliary elements in the larger fragment in ars2-3358. The presence of auxiliary sequences and their possible role in enhancing origin activity reported previously for *ars2004* [15]. Most of the ars2-3358(2452M) maintained as multimers in *S. pombe* cells (as assayed in Brun et al. (1995))(Table 2), a property of plasmids containing weak origins, reflecting the genomic efficiency of this origin [20].

**Table 2.** Consolidated table enlisting all plasmids, their size, AT%, relative transformation frequency mitotic stability and maintenance status of plasmids

| Name of Plasmids | Fragment size (kb) | AT% | ARS +/- | Relative transformation frequency | Mitotic stability (%) | Maintenance status of plasmid |
| --- | --- | --- | --- | --- | --- | --- |
| p3002 | 1.2 | 71.4 | +ve | 1 | 76.8 | .... |
| ars2-3335 | 1.5 | 68.9 | +ve | 0.64 | 54.6 |  |
| ars2-3358(2452M) | 2.4 | 76.3 | +ve | 0.32 | 38.5 | Multimer |
| ars2-3363 | 2.3 | 68.7 | +ve | 0.58 | 55.8 | Monomer/<br>dimer |
| ars2-3363pd | 2.1 | 69.1 | +ve | 0.35 | 52.34 | .... |
| ars2-3374 | 2.4 | 65.5 | -ve | 0.004 | 15.5 | Dimer/<br>Multimer |
| ars2-3398 | 2.3 | 67.3 | +ve | 0.24 | 41.4 | Dimer, Multimer |
| p2482 | 2.5 | 67.1 | -ve | 0.003 | 14.9 | Multimer |

The II-3363 containing plasmid ars2-3363 maintains its status as a monomer (Table 2) and is capable of transforming cells at a higher frequency, reflecting the genomic origin efficiency reported earlier [20]. It has two transcriptional promoters at its right end. Promoters present near origins have been reported to enhance origin activity of replicators in yeasts [34,35]. To see how the ars2-3363-associated promoters influence its origin activity, we deleted these promoters by PCR based cloning resulting into ars2-3363pd. The transformation efficiency of ars2-3363pd is consistently slightly lower than ars2-3363 with a little effect on its mitotic stability (Figure 1C-E). These differences are not statistically significant and probably reflect the effect of the size reduction as seen in case of ars2-3358(2452M). These results corroborate an earlier report from Gómez & Antequera (1999), in which they showed that deletion of a 231-bp transcription initiation region abolishes the transcription but has little effect on the replication of *ars1* of *S. pombe*.

A comparison of the transformation frequencies and mitotic stabilities of the plasmids derived from the five 2D mapped origins shows that the plasmids with the three moderate origins (II-3335, II-3363 and II-3398), ars2-3335, ars2-3363 and ars2-3398 are more stable than those derived from weak origins (II-3358 and II-3374), ars2-3358 and ars2-3374 reflecting their chromosomal origin efficiency (Figure1F, Table 2). Our origin efficiency estimates for these origins are very similar to that reported by Daigaku et al. (2015) who found 9 origins in this region using a replicative polymerase profiling method (Figure 1F). It is likely that the cloned DNA fragments contain most, or all essential genetic information required for the activity of these origins.

### Sequence analysis of ARSs

Next, we analyzed sequences of the ARSs to identify similarities between them and to identify any motifs or common sequence characteristics. To identify AT-rich motifs, which we call polyATs in this study, in these sequences, we searched for 10 or more contiguous As or Ts., because sequences of ARSs are highly AT-rich (Table 2) [10,16,20,26]. We found polyAT motifs in all 5 ARSs as well as in non-origin fragment (p2482) (Table 3). We used BLAST to look for the occurrence and distribution, or near matches occurrence and distribution of these polyATs in the genome (see Material and Methods for details). We found over ∼2300 near matches of polyAT motifs across the genome (Supplementary data 1 and Table 4). Interestingly, none of the motifs from ARS negative fragment (p2482) or week origin (ars2-3374) showed near matches in the genomic BLAST. We then compared these near match polyATs within ARS (e.g., ars2-3335 vs ars2-3335), with other ARSs (ars2-3335, ars2-3358, ars2-3363, and ars2-3398 (Table 5) and vice versa, to find out similarities among the polyATs if any. However, very few of them were overlapped (Supplementary data 1, Table 5) with each other, which shows these are unique sequence motifs. Moreover, we looked for a consensus motif in 2309 polyATs, (Supplementary data1), using the online MEME motif identification tool (http://meme-suite.org/tools/meme, Figure 2G (A/TA/TA/TA/TAAA/TA/TAA/GorCA/G).

**Table 3:**
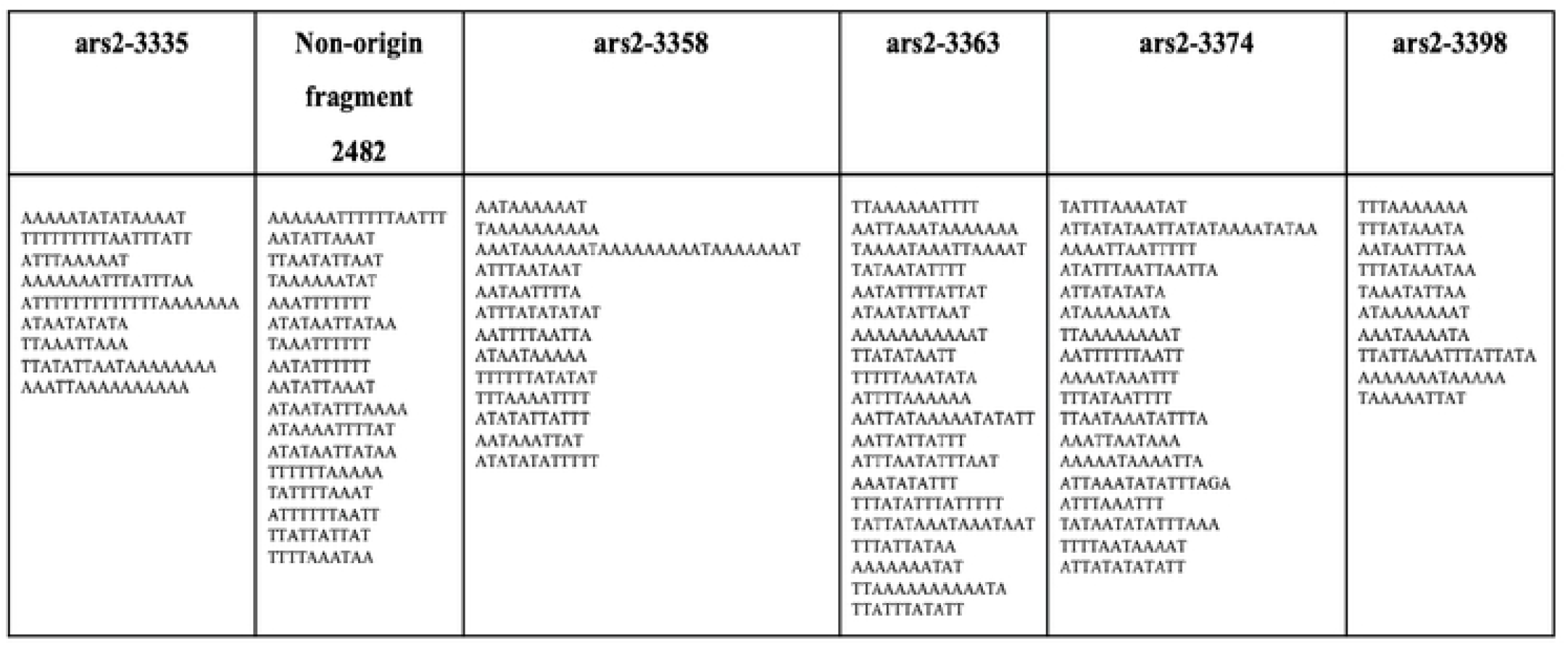
PolyAT sequences found in all five ARSs and non-origin fragment.

**Table 4:**
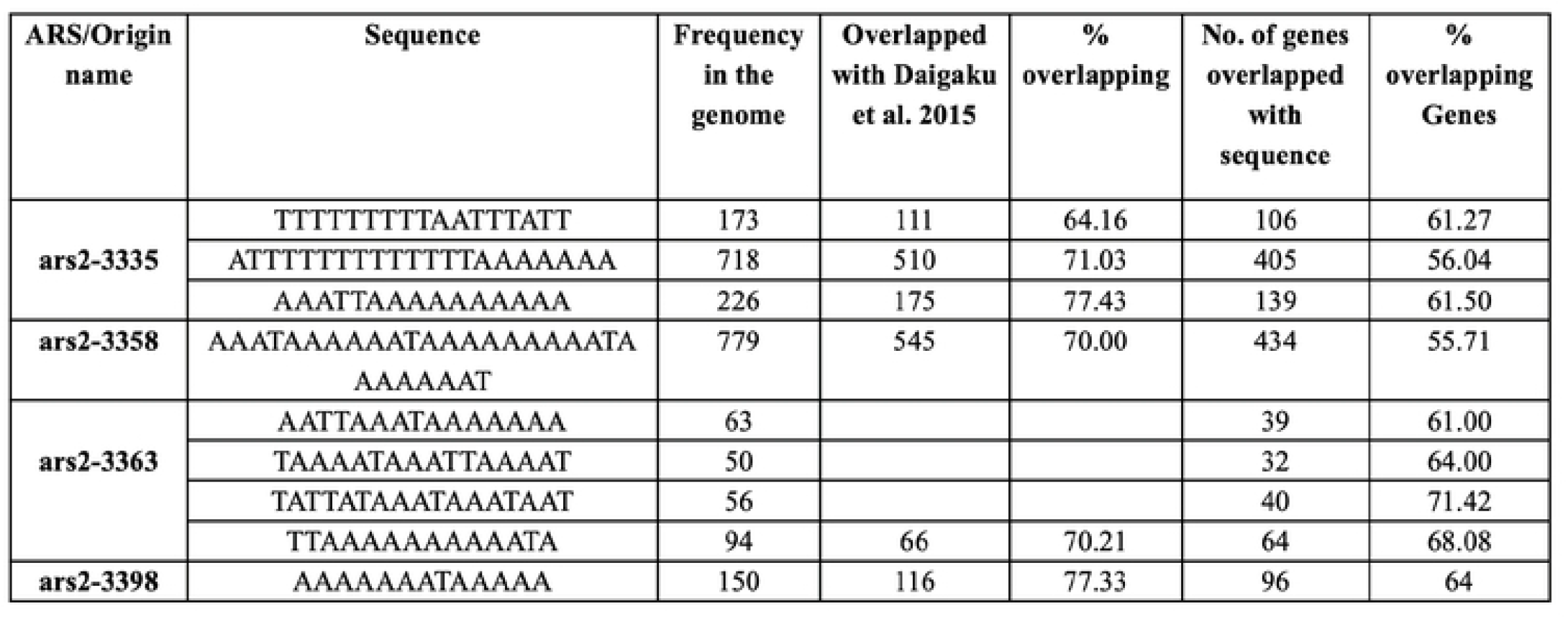
PolyAT sequences and their near match frequencies in the genomic BLAST search, % overlapping with genome-wide origins and genes in the genome

**Table 5:**
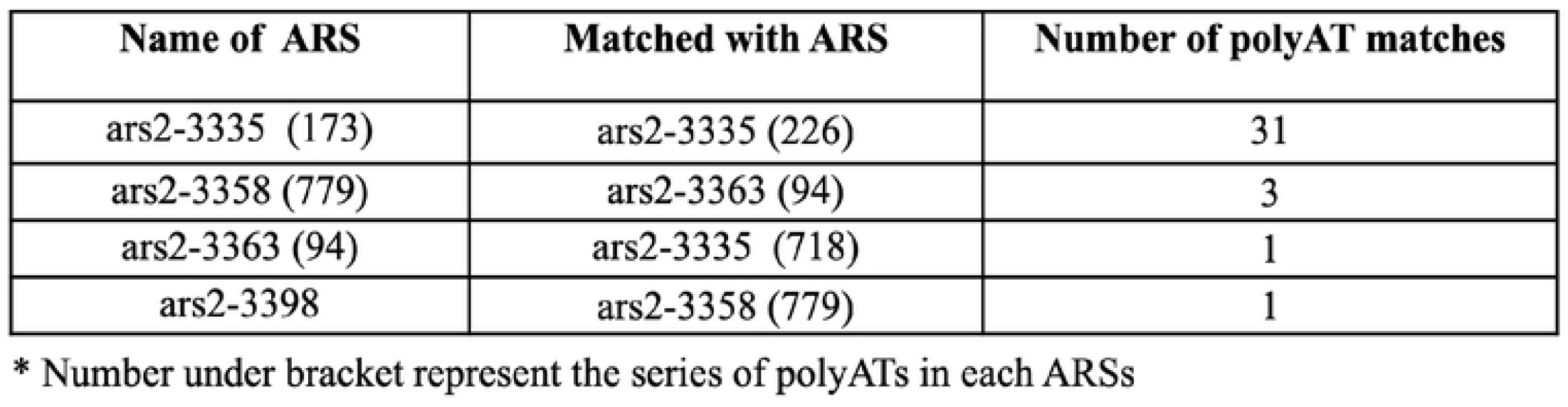
PolyAT sequence matches within ARSs and with other ARSs.

**Figure 2:**
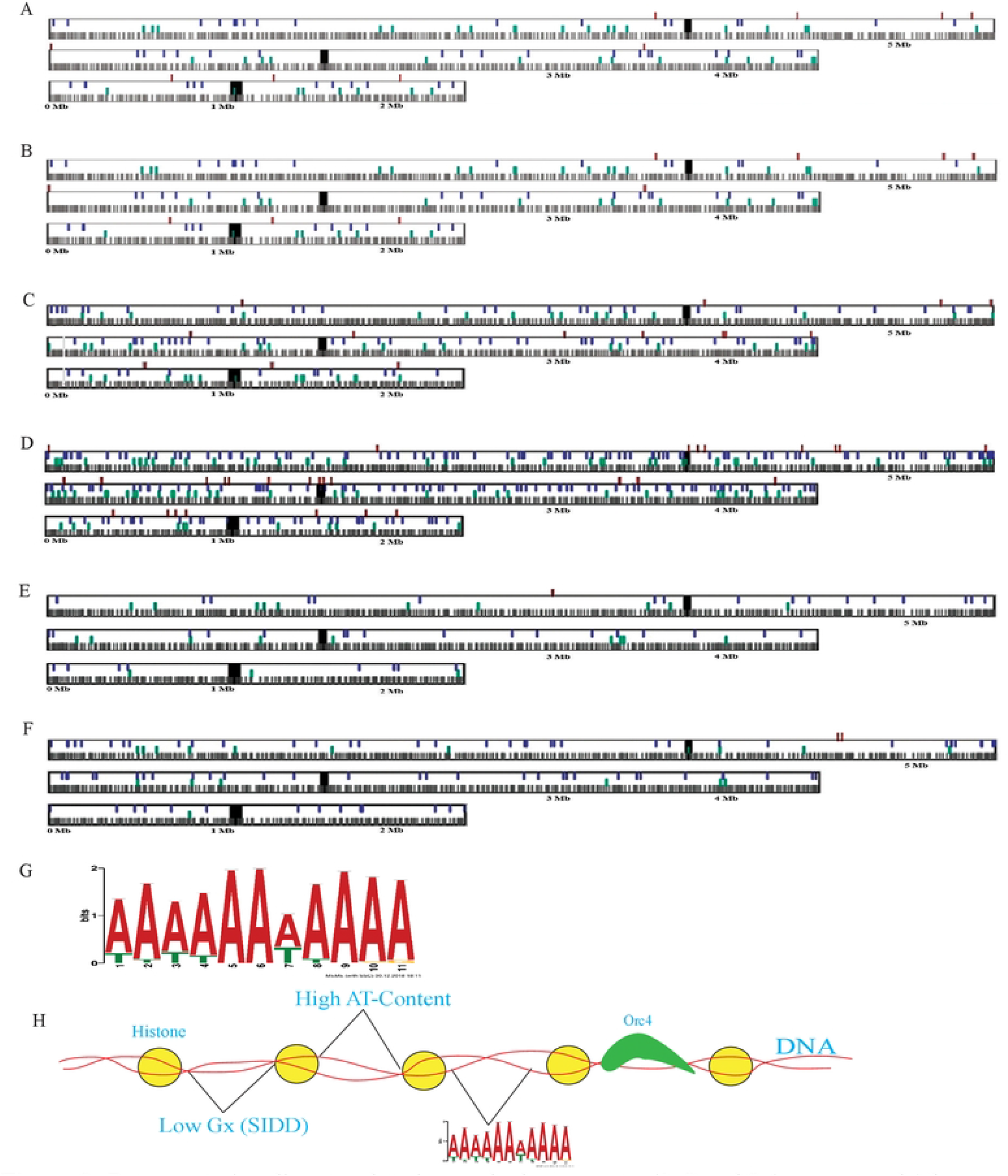
Representative diagram showing each chromosome (1, 2 and 3 (as a rectangle) known origins of Daigaku et al. 2015 (black verticle thin lines) overlaps with polyATs (red, green and blue) (and thick black line showing centromere in each chromosome. (A-C) ars2-3335 polyAT near match -173, 226 and 718 (TTTTTTTTTAATTTATT, AATTAAAAAA AAAA and ATTTTTTTTTTTTTA -AAAAAA), (D) ars2-3358 polyW near match 779 (AAATAAAAAATAAAAAAAAATAAAAAAAT) (E) ars2-3363 near match 94 (TTAAAAAAAAAATA), (F) ars2-3398 poly AT near match 150(AAAA -AAATAAAAA), (G) 11-bp Motif, found using MEME motif search online tool and (H) hypothetical model for DNA replication.

Next we compared polyATs positions with known origin positions [21] taking 1-kb upstream and downstream, >65% of the polyATs (∼2300) overlapped with known origins (Supplementary 1, Table 4, Figure 2A-F) [21]. The average distribution per kb of polyATs is 19-25-kb, which coincides with the an earlier estimated frequency of one ARS per 20-50-kb [9,37,38] (Supplementary data1; Figure 2A-F). However, twelve thousand ACSs had been identified in the genome of *S. cerevisiae* though only 400 of these (3.3%) were found to be functional [39,40].

We have also tested the functional importance of polyATs published in earlier studies to determine whether they modulate ARS activity or not. Systematic comparison with few deletion, mutation and substitution studies done earlier, found significant overlap with each of the study [10,14–16,18,41], showing clear modulation in the efficiency of ARSs or origins (Supplementary Figure 1).

The polyAT motifs found during this study overlap well with the *ars1, ars3001* and *ars3002* sequences found in the functional and interactor studies to identify interactors ORC4 of *S. pombe* [23,42–45]. The origin recognition complex (ORC) recognizes origins by the binding of AT-hooks present at the N-terminal end of ORC4 to ∼4-6 bp AT-rich sequences and the AT-hooks are essential for ORC assembly to initiate replication at *S. pombe* origins [23,42–45]. Our study reflects the importance of polyATs found in this study and could be used as a *S. pombe* origin predictor [46]. However, with all overlaps and importance of the polyAT for origin activity, we propose a hypothetical model to summarize our study, *S. pombe* origins having broader nucleosome free regions with a polyAT motif to bind ORC4 and coincide with lowest SIDD (Figure 2H and Figure 3A-E).

**Figure 3:**
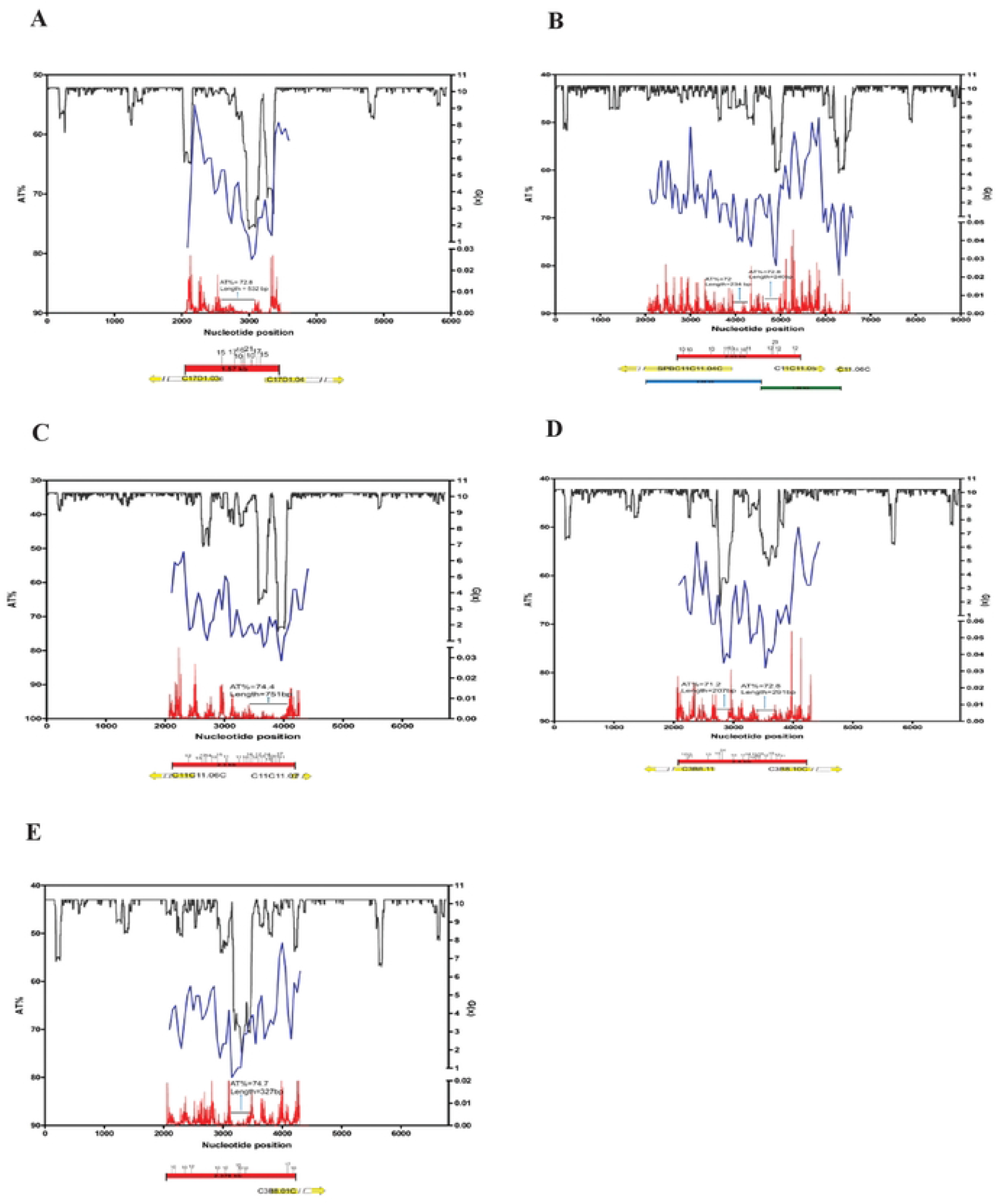
SIDD profiles, nucleosome occupancy and AT% of 50-bp overlap of the cloned fragment in pRS306 background calculated by WEBSIDD (black line), Segal’s lab (Kaplan et al. 2008; http://genie.weizman.ac.il/software/nucleo_prediction.html) (redline) and AT% 5-bp overlap (blue line(analysis done using FastPCR). The locations of the genes (yellow arrows) flanking the cloned fragment (red thick line) is marked with stretches of polyATs (small black vertical lines with number indicating length of polyAT stretches). (A) ars2-3335, (B)- ars2-3358, (C) ars2-3363, (D) ars2-3374 and (E) ars2-3394

## Conclusion

Our study elucidates a role of polyATs in origin recognition and suggests the importance of polyATs in origin efficiency. The majority (>65% to >77%) of polyATs overlap with genome-wide origins pave the way for the future study to focus on their functional role in origin activity.

## Acknowledgements

This work was supported by the Council of Scientific and Industrial Research (CSIR) -New Delhi by providing Senior Research Fellowship to M.P.Y. ref. no. 9/1014(0002)2k11-EMR-I and University Grant Commission (UGC)-New Delhi by providing research grant to Prof. D.D. Dubey, ref. no.-32-551/2006(SR). All the research reported in this publication included work performed in Department of Biotechnology, Veer Bahadur Singh, University Jaunpur, Uttar Pradesh-India. We are very thankful to late Dr. Vinay Kumar Srivastava for giving few plasmids for the study. We would like to heartly thank Prof. Nick Rhind for reading this manuscript, critics, suggestions and correction of manuscript, which are greatly appreciated by the authors.

## Author contributions

M.P.Y. and D.D.D. contributed to project planning and designing the experiments. M.P.Y. performed all the experiments and wrote the manuscript.

## Competing interests

Authors declare no competing interests.

## Notes

### Competing Interest Statement

The authors have declared no competing interest.

